# Alpha modulation of spiking activity across multiple brain regions in mice performing a tactile selective detection task

**DOI:** 10.1101/2025.03.07.642074

**Authors:** Craig Kelley, Cody Slater, Marc Sorrentino, Dillon Noone, Jocelyn Hung, Paul Sajda, Qi Wang

## Abstract

Many cognitive and sensory processes are characterized by strong relationships between the timing of neuronal spiking and the phase of ongoing local field potential oscillations. The coupling of neuronal spiking in neocortex to the phase of alpha oscillations (8-12 Hz) has been well studied in nonhuman primates but remains largely unexplored in other mammals. How this alpha modulation of spiking differs between brain areas and cell types, as well as its role in sensory processing and decision making, are not well understood. We used Neuropixels 1.0 probes to chronically record neural activity from somatosensory cortex, prefrontal cortex, striatum, and amygdala in mice performing a whisker-based selective detection task. We observed strong spontaneous alpha modulation of single-neuron spiking activity during inter-trial intervals while mice performed the task. The prevalence and strength of alpha phase modulation differed significantly across regions and between cell types. Phase modulated neurons exhibited stronger responses to both go and no-go stimuli, as well as stronger motor- and reward-related changes in firing rate, than their unmodulated counterparts. The increased responsiveness of phase modulated neurons suggests they are innervated by more diverse populations. Alpha modulation of neuronal spiking during baseline activity also correlated with task performance. In particular, many neurons exhibited strong alpha modulation before correct trials, but not before incorrect trials. These data suggest that dysregulation of spiking activity with respect to alpha oscillations may characterize lapses in attention.

## Introduction

Neural oscillations play a critical role in regulating various perceptual and cognitive functions, including sensory processing, memory, attention, and decision-making. They serve as a mechanism for coordinating communication between different neural circuits (Gray et al., 1989; Buzsáki and Draguhn, 2004; Weiss et al., 2023). Since the advent of electroencephalography (EEG), alpha oscillations (7-12 Hz, ranges vary depending on the study) have been the object of close study (Berger, 1929; Ojha, 2024). Alpha oscillations have been observed in diverse cortical regions, including visual, somatosensory, motor, premotor, auditory, and entorhinal cortices (da Silva et al., 1973; Steriade et al., 1990; Lehtelä et al., 1997; Lukatch and MacIver, 1997; Castro-Alamancos and Tawara-Hirata, 2007; Haegens et al., 2011). Because alpha oscillations are most prominent in visual cortex with eyes closed, they were previously hypothesized to represent an idling state and were therefore inessential for understanding cognition (Adrian and Matthews, 1934; Steriade et al., 1990; Worden et al., 2000). However, there is strong evidence that alpha oscillations play a functional role in cognitive processing with rhythmic inhibition suppressing the representation and processing of task-irrelevant stimuli (Cooper et al., 2003; Klimesch et al., 2007; Palva and Palva, 2007). Lateralization of alpha oscillations in spatial attention tasks, with high alpha power in task-relevant regions ipsilateral to attended stimuli (in the unaligned, non-encoding hemisphere relative to the stimuli) and low alpha power in task-relevant regions contralateral to attended stimuli (in the aligned, encoding hemisphere relative to the stimuli), provide further support for the role of alpha in suppressing irrelevant representations (Jensen and Mazaheri, 2010; Händel et al., 2011; Wildegger et al., 2017; Dahl et al., 2019; Schneider et al., 2019; Bagherzadeh et al., 2020). This line of evidence, however, might suggest that alpha oscillations are unimportant in the brain hemisphere contralateral to (aligned with) behaviorally relevant stimuli. Several studies have suggested that alpha rhythms represent a preparatory brain state, that is prepared to receive and respond to sensory stimuli (Adrian and Matthews, 1934; Nikouline et al., 2000; Fanselow et al., 2001; Cooper et al., 2003; Tort et al., 2010b). The rhythmic inhibition associated with alpha oscillations may provide a mechanism by which sensory processing is discretized (VanRullen and Koch, 2003; Mazaheri and Jensen, 2010). In the present study, we focused on alpha band oscillations in the hemisphere aligned to attended stimuli.

Much of the work on alpha oscillations’ role in human cognition has focused on changes in power or synchronization in the EEG over long timescales (on the order of seconds), without accounting for the relationship between neuronal firing and the alpha rhythm on the timescale of an alpha cycle (∼80 - 125 ms). Several studies have demonstrated phase locking of neuronal firing to cortical alpha oscillations in nonhuman primates, however, demonstrating coupling of cortical spiking activity to the phase of the alpha cycle in local field potentials (LFPs) (Bollimunta et al., 2008; Bollimunta et al., 2011; Buffalo et al., 2011; Haegens et al., 2011; van Kerkoerle et al., 2014; Dougherty et al., 2017). Cortical firing activity was shown to be elevated at the troughs of alpha oscillations and decreased during the peaks, supporting the rhythmic inhibition hypothesis (Haegens et al., 2011). The origins of alpha oscillations, and the identities of the neurons that serve as pacemakers for that rhythm, vary by cortical region (Lukatch and MacIver, 1997; Jones et al., 2000; Bollimunta et al., 2008; Bollimunta et al., 2011). Alpha phase modulation of spiking activity in the primary visual cortex has been shown to regulate visually driven responses (Dougherty et al., 2017). Other studies have investigated the phase locking of cortical spiking activity to alpha, other oscillatory bands in the LFP, or to a “generalized” LFP phase (5 - 50 Hz) and showed differences in the prevalence and strength of phase modulation change with cortical layer (Lakatos et al., 2005; Bollimunta et al., 2008; Davis et al., 2023).

Despite the extensive work on alpha modulation of cortical spiking activity in nonhuman primates, the phenomenon remains largely unexplored in other mammals and brain regions. Additionally, most studies of alpha phase modulation in nonhuman primates focused on evoked multi-unit activity, without distinguishing which cell types were coupled to the alpha oscillation (Bollimunta et al., 2008; Bollimunta et al., 2011; Buffalo et al., 2011; Haegens et al., 2011; van Kerkoerle et al., 2014; Dougherty et al., 2017). We therefore used Neuropixels 1.0 probes to record LFPs and single-unit spiking activity in task-relevant brain regions of mice as they performed a whisker-based selective detection task based on Aruljothi et al. (2020) and quantified the modulation of single neurons’ baseline spiking activity by the phase of alpha oscillations. We found differences in the prevalence and strength of alpha phase modulation between brain regions (somatosensory cortex, prefrontal cortex, striatum, and amygdala), between cell types (regular spiking and fast spiking units) within those regions, and along dorsoventral axes within cortical regions. We found that phase modulated neurons exhibited greater task-related changes in firing rate compared to neurons that were not phase modulated, and we explored how phase modulation of spiking activity mapped onto task performance. Our results suggest that alpha phase modulation of neuronal firing during spontaneous activity promotes representation of, and correct responses to, behaviorally relevant stimuli.

## Methods

All experimental procedures were approved by the Columbia University Institutional Animal Care and Use Committee (IACUC protocol number: AC-AABN8555) and were conducted in compliance with NIH guidelines. Adult mice of both sexes (3 females, 1 male), aged 3 - 12 months, were used in the experiments. All mice were kept under a 12-hour light-dark cycle. All software used for controlling the behavioral apparatus and data analysis can be found at https://github.com/Neural-Control-Engineering/n-CORTex which contains project-specific code, and https://github.com/Neural-Control-Engineering/AlphaModulation_SelectiveDetection which provides general purpose tools for neurophysiological and behavioral data collection and segmentation.

### Behavioral task

Head-fixed mice (n=4) were trained to perform a whisker-based go/no-go selective detection task based on Aruljothi et al. (2020). During behavioral training mice were water restricted to maintain ≥ 85% of their post-surgery body weight. Mice had ad libitum access to food. The behavioral apparatus was controlled by custom code written in MATLAB and Simulink (MathWorks, Natick, MA) via a Speedgoat real-time system (Speedgoat Inc., Natick, MA). Trials began with a unilateral whisker deflection lasting 200 ms delivered via a piezoelectric actuator connected to a flat paddle which made contact with multiple whiskers while the mouse was not actively whisking. The paddles were not physically attached to the whiskers, so the mice could freely whisk over the course of the session. The tactile stimulus was driven by a triangle wave resulting in a 1 mm deflection of the paddle with 100 ms of rise time and 100 ms of fall time (**Fig. 1**). Mice were required to selectively respond to left-sided whisker deflections (go stimuli) by licking a water spout within a 1 s window of opportunity in order to receive a water reward, and they were required to ignore right-sided whisker deflections (no-go stimuli). Failure to withhold licks during the window of opportunity following a no-go stimulus resulted in an increased inter-trial interval (8-12 s with a negative exponential distribution, as opposed to 4-8 s otherwise). On any given trial, the probability of a go stimulus presentation was 30%; the probability of a no-go stimulus was 70%. Mice had to withhold licks during the 200 ms of stimulus delivery. Licking during this period voided the trial (it was not included in further analyses, and mice did not receive water reward regardless of the side on which the stimulus was delivered). Licking was allowed during the intertrial period, however. Sessions lasted one hour. All mice achieved expert performance on the task, defined as 3 consecutive sessions with perceptual sensitivity (d’) greater than 1, and we included sessions after and including the first session in that 3-session streak (**Fig 1D**). The lateralization of the go and no-go stimuli remained fixed across sessions.

**Figure 1:**
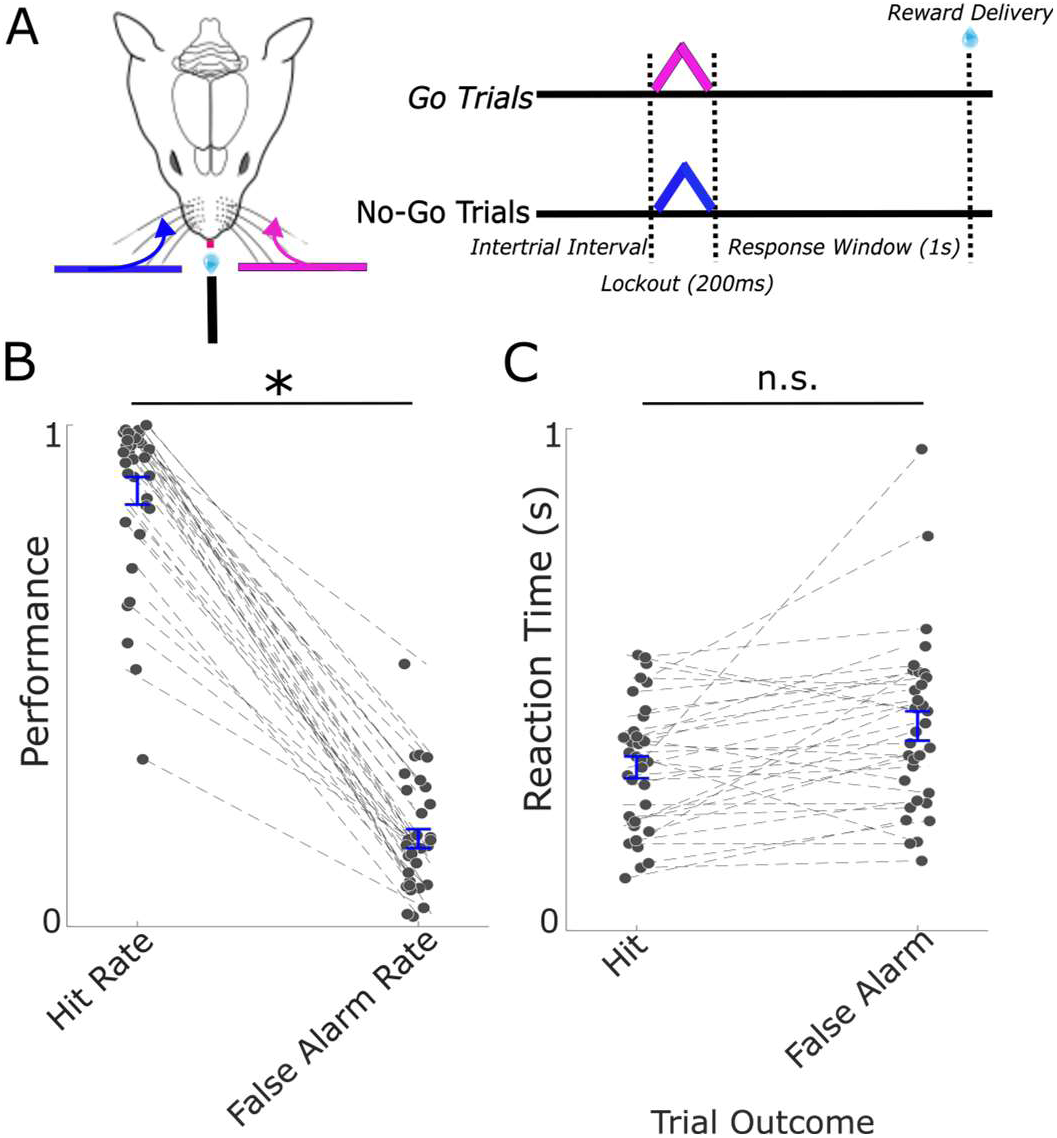
Mice expertly performed a whisker-based selective detection task. **(A)** Schematic representation of a whisker-based selective detection task. Head-fixed mice were required to lick a waterspout during a 1 s response window following left-sided whisker deflections (go stimuli) to receive a water reward. They were required to withhold licks in response to right-sided deflections (no-go stimuli) during the window of opportunity to avoid a longer inter-trial interval. **(B)** Expert mice exhibited significantly higher hit rates than false alarm rates. Average hit and false alarm rates shown in blue. **(C)** Reaction times did not differ significantly between hits and false alarms. Data from individual sessions are connected by dashed lines. Average reaction times shown in blue. * indicates p < 0.05.

### Surgical procedures

Prior to behavioral training, mice were implanted with a Neuropixels 1.0 electrode and head bar. Animals were anesthetized with isoflurane in oxygen (5% induction, 1-2% maintenance) and fixed in a stereotaxic frame (David Kopf Instruments, CA). Body temperature was maintained at 37℃ using a feedback-controlled heating pad (FHC, Bowdoinham, ME). When the animal’s condition stabilized, buprenorphine (0.05 mg/kg) was administered subcutaneously before an incision on the scalp was made. Each mouse was implanted with a Neuropixels 1.0 multielectrode encased in a custom built headstage housing designed for chronic head-fixed recordings (Liu et al., 2024) (**Fig. 2A**). Probe trajectories were planned and monitored in real-time during implantation using Neuropixels Trajectory Explorer (https://github.com/petersaj/neuropixels_trajectory_explorer, by Andy Peters). Two mice were implanted with probes spanning primary somatosensory cortex, striatum (dorsal striatum / caudoputamen and small portions of ventral striatum), and basolateral amygdala contralateral to the side where go stimuli were presented (see Behavioral task below) (**Fig. 2A, B**). Two other mice were implanted with a probe in medial prefrontal cortex (primarily infragranular layers of anterior cingulate, pre- and infralimbic, orbitomedial, and dorsal peduncular cortices) (**Fig. 2B**) contralateral to the go stimulus. Baytril (5 mg/kg) and Carprofen (5 mg/kg) were administered at the end of the surgery and every 24 hours after the surgery for four days. Animals’ weight was measured once a day during this post-operative period. Aruljothi et al. (2020)

**Figure 2:**
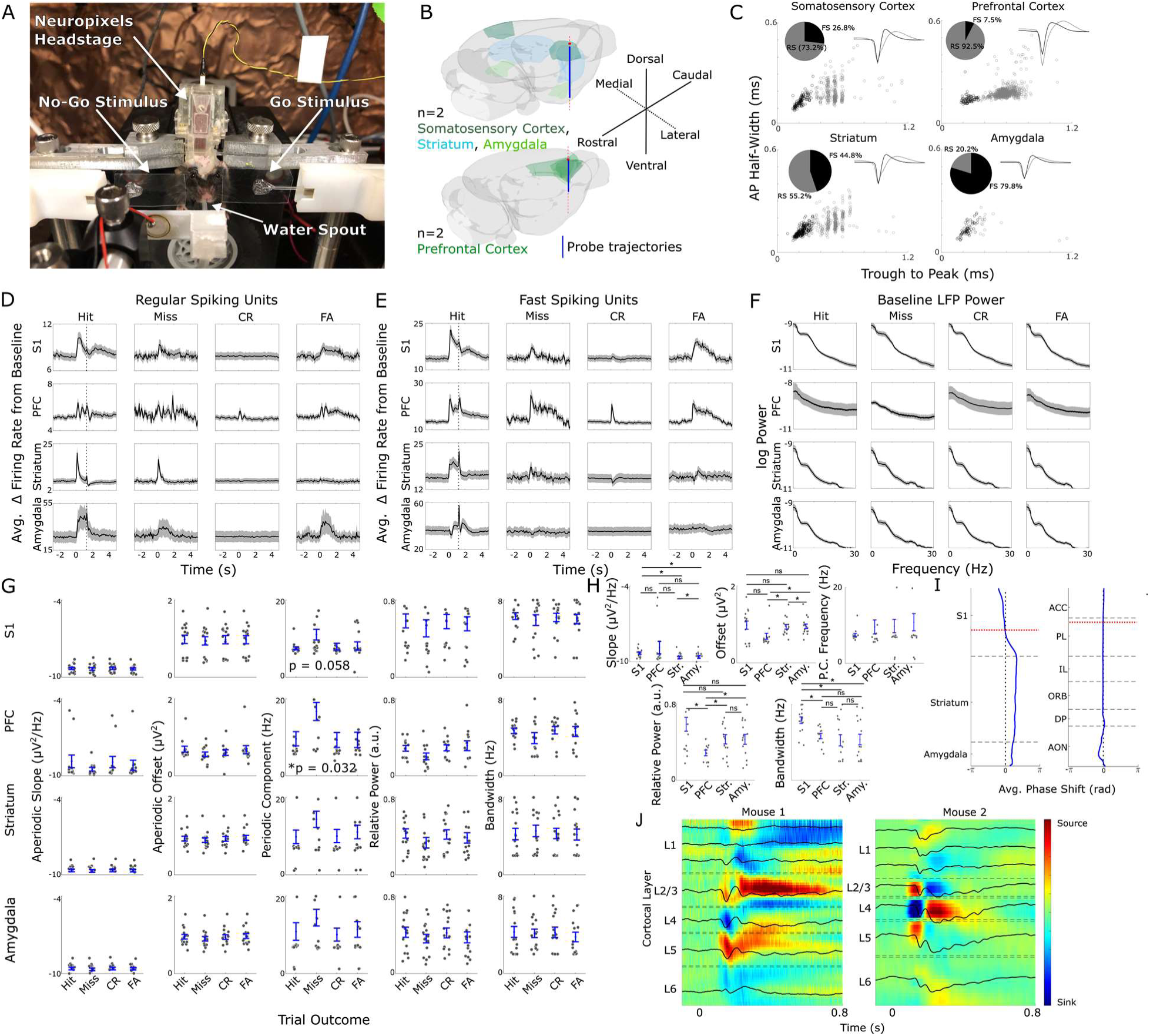
Neural activity across different brain regions exhibited task-related dynamics during the tactile selective detection task. **(A)** Photo of head-fixed mouse with chronically implanted Neuropixels 1.0 electrode in S1, striatum, and amygdala performing the whisker-based selective detection task. **(B)** 3D renderings of probe trajectories planned with Neuropixels Trajectory Explorer. **(C)** Classification of RS and FS units in S1, PFC, striatum, and amygdala based on the time from the trough of the waveform to the subsequent peak and the width of the negative deflection. **(D)** Average firing rates of all RS units in S1 (n = 325), PFC (n = 769), striatum (n = 334) and amygdala (n = 17) aligned to the tactile stimulus and separated by trial outcome. **(E)** Average firing rates of all FS units in S1 (n = 120), PFC (n = 50, center), striatum (n = 271), and amygdala (n = 66) aligned to the tactile stimulus and separated by trial outcome. **(F)** Average LFP spectra separated by trial outcome across all sessions from electrodes in S1 (layer 4), PFC (PL), striatum (caudoputamen), and amygdala (lateral amygdala). **(G)** Summary of aperiodic (slope and offset) and periodic components (local peak center frequencies, relative power, and bandwidth) of LFP spectra used to compute averages shown in panel **F**. **(H)** Inter-regional comparisons of aperiodic and periodic LFP components. **(I)** Average alpha-band phase shift across brain regions relative to reference electrodes (represented with white lines; left: S1 layer 4; right: PL). **(J)** CSD in S1 average across sessions for each mouse with an implant spanning S1, striatum, and amygdala. * indicates p < 0.05. In panels without p-values or *, there were no significant differences.

### Neural recordings

Neural signals across 384 electrodes of the Neuropixels 1.0 probe were recorded while the mice performed the selective detection task using a National Instruments PXI system controlled by SpikeGLX (Release v20230905-phase30). The recording system was synchronized with the behavioral apparatus via TTL pulses generated by the Speedgoat real-time system. The behavioral task and neural recordings were performed in a box lined with copper mesh to reduce electromagnetic noise. SpikeGLX wrote data to disk in two separate binary files: one containing a wide-band signal (0.3-10 kHz bandwidth, 30 kHz sampling rate, 250x gain) and the other a lower band signal (0.5-500 Hz bandwidth, 2.5 kHz sampling rate, 125x gain) for analysis of the local field potential (LFP). Single-unit spiking activity was extracted offline using Kilosort 4.0 (Pachitariu et al., 2024). Only single-units labelled ’good’ by Kilosort 4.0 were used in further analysis. All spike waveform templates and single-unit autocorrelograms were visually inspected, and neurons with distorted waveform templates, physiologically unrealistic inter-spike intervals, or artifacts related to line noise in their spiking activity were rejected from further analysis. Local field potentials (LFPs) were downsampled to 500 Hz after applying a fourth order Butterworth bandpass anti-aliasing filter with cutoff frequencies at 0.1 Hz and 100 Hz using forward and backward passes (MATLAB’s *filtfilt)*.

### Analysis

Individual neurons were classified by their waveform templates identified by Kilosort 4.0. Positive spiking units, characterized by positive deflections in the extracellular waveform and likely representing axonal spikes, and triphasic spiking units, characterized by small positive deflections preceding larger negative deflections and likely representing dendritic spikes, were not included in further analyses (Jung et al., 2023; Someck et al., 2023). The remaining units were classified as regular spiking (RS) and fast spiking (FS) units based on the action potential width and the time between the trough and subsequent peak of the waveform template using k-means clustering (**Fig. 2C**) (Hocker et al., 2021; Liu et al., 2024). Peristimulus time histograms and firing rates were computed using 50 ms non-overlapping bins. Baseline multitaper LFP spectra (time-bandwidth product 5, 9 tapers) were computed using the Chronux toolbox (http://chronux.org/) (Mitra and Bokil, 2007). We also analyzed the aperiodic (1/f-like dynamics) and periodic (local peaks in the spectrum) components of averaged spectrograms using an algorithm developed by Donoghue et al. (2020) (https://github.com/fooof-tools/fooof). For each session, we averaged baseline LFP spectra across trials of each outcome, identified local peaks in the averaged spectra, and computed the relative power (to the rest of the spectrum) and bandwidth of those peaks.

To identify neurons whose baseline activity was significantly modulated by alpha phase, we first isolated alpha band activity in the LFP on each electrode where a single-unit was recorded by bandpass filtering the raw LFP signal from 8 to 12 Hz with a fourth order non-causal Butterworth filter (MATLAB’s *filtfilt)* to avoid imposing spurious phase shifts. The instantaneous phase of the alpha band was computed using the Hilbert transform. Single-unit spikes were then registered to the instantaneous phase of alpha on the electrode where the spike waveform produced the greatest deflection. We then pooled the instantaneous phases of all baseline (3 s before stimulus) spikes across the session and used Rayleigh’s test for non-uniformity (Fisher, 1993). Units were considered significantly phase modulated if the p-value from Rayleigh’s test was less than 0.05 divided by the number of units recorded on a given session (Bonferroni correction for multiple comparisons). To quantify alpha modulation of spiking activity, we fit a von Mises distribution (the circular analog of normal distribution) to the probability density function of spikes in 20 18° bins of alpha phase (Eschenko et al., 2012). A similar procedure was used to identify whether units were phase-modulated prior to trials of various outcomes, only including instantaneous phases of baseline spikes on trials of a given outcome. The same corrected p-value threshold was used to identify significant phase modulation prior to trials with different outcomes. This same procedure was also used to compare phase modulation on trials with spontaneous licking (prior to the stimulus presentation) to phase modulation on trials with no spontaneous licking. Comparisons of preferred phase (phase of alpha at which the neuron is most likely to produce an action potential) between cell types and regions were performed using Kuiper’s test (Fisher, 1993). We computed the modulation index (MI), which measures how the observed distributions of spikes relative to alpha phase deviated from a uniform distribution, to quantify how the strength of alpha modulation (Tort et al., 2010a; Dougherty et al., 2017):

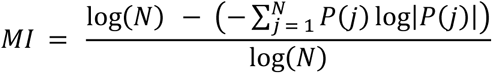

Here, *j* is a single phase bin and *P(j)* is the normalized spike count in that bin. For the handful of neurons that exhibited bimodal distributions of spiking probability relative to alpha phase, we fit a mixture model of two von Mises distributions.

We also compared alpha phase modulation during oscillatory events with high and low alpha power during spontaneous activity (3 s). For each phase modulated neuron, we computed instantaneous alpha power on the electrode where that neuron was recorded using bandpass filtering and the Hilbert transform as above. We identified epochs 0.3 s or longer where alpha power was greater than or equal to the 75th percentile of alpha power across the session on that electrode as high alpha events and epochs 0.3 s or longer where alpha power was less than the 50th percentile of alpha power across the session as a low alpha event. When there was an interval between events of the same kind lasting 0.2 s or shorter, those events were combined. We then analyzed phase modulation of spiking activity across high alpha power events and low alpha power events separately.

### Statistics

All circular statistics were performed using the CircStat toolbox for MATLAB (Berens 2009). We used Rayleigh’s test for non-uniformity to determine which neurons were significantly modulated by alpha phase. We used Kuiper’s test to compare pairs of distributions of preferred phase angles (the alpha phase at which a neuron is most likely to spike) (Fisher, 1993). Watson-Williams test was used to compare preferred phase angles across cortical layers and among prefrontal subregions (Berens 2009). For non-circular data, we tested for homoscedasticity using the Kolmogorov-Smirnov test. If data were not normally distributed, we used nonparametric tests (Wilcoxon signed-rank test and Mann-Whitney U-test); otherwise, we employed parametric tests (paired and two-sample t-test) with one exception. To compare percentages of responsive neurons between alpha modulated units and unmodulated units, we employed Fisher’s exact test. Data are presented as mean ± standard error unless noted otherwise.

## Results

We trained head-fixed mice (n=4) to perform a whisker-based selective detection task, requiring mice lick in response to left-sided whisker deflections (go stimuli) to receive a water reward and withhold licks in response to right-sided whisker deflection (no-go stimuli) to avoid a longer intertrial interval (**Fig. 1A**). Performance was evaluated using discriminability (d’) from signal detection theory (Stanislaw and Todorov, 1999). The average d’ across all sessions was 2.3 ± 0.14, with hit rates (87.0 ± 2.7%) significantly higher than false alarm rates (17.5 ± 1.9%) (**Fig. 1B**; p = 4.6 x 10^-23^, paired t-test). Reaction times did not differ significantly between false alarm trials (0.50 ± 0.05 s) and hit trials (0.64 ± 0.15 s; p = 0.60, paired t-test; **Fig. 1C**).

We recorded neural activity during expert task performance across 34 sessions with chronically implanted Neuropixels 1.0 probes (**Fig. 2A**). Probe trajectories spanned primary somatosensory/barrel cortex (S1), striatum (mostly dorsal striatum / caudoputamen), and amygdala (lateral and basolateral amygdala) in 2 mice and prefrontal cortex (PFC) in 2 other mice (**Fig. 2B**). Single-units were classified as regular spiking (RS; mostly putative excitatory neurons) and fast spiking (FS; mostly putative inhibitory neurons). Of the 356 units recorded in S1, 73.2% were classified as RS units and 26.8% of units were classified as FS units. Of the 442 units recorded in PFC, 92.5% were classified as RS units and 7.5% of units were classified as FS units. Proportions of FS and RS units recorded were in line with previous observations in S1 and PFC (Agmon and Connors, 1992; Liu et al., 2024). Of the 328 units recorded in striatum, 55.2% of units were classified as RS units, while 44.8% were classified as FS units. This was a more equal distribution than previous observations from striatum in rats and nonhuman primates (Gage et al., 2010; Yamada et al., 2016). Finally, of the 74 units recorded in amygdala, 20.2% were classified as RS and 79.8% were classified as FS. Previous work found a higher percentage of FS units in amygdala in rats but used firing rate rather than waveform shape to classify units (Rosenkranz and Grace, 1999).

Subpopulations in each recorded region exhibited task-related spiking dynamics during task execution and differences in stimulus-evoked changes in spiking activity during the 200 ms decision period between stimulus presentation and acceptable motor response (**Fig. 2D, E**). We focused on comparisons between hit and miss trials because neural recordings were aligned (contralateral) to the go-stimulus. FS units in S1 showed significantly greater stimulus-evoked increases in firing rate from baseline during hit trials (11.0 ± 1.2 Hz) than on miss trials (7.4 ± 1.1 Hz; p = 2.9x 10^-6^, Wilcoxon signed-rank test), while there was no significant difference in stimulus-evoked activity in S1 RS units (3.5 ± 0.3 Hz on hit trials, 2.5 ± 0.3 Hz on miss trials, p = 0.10, Wilcoxon signed-rank test). In PFC, neither RS nor FS units exhibited significantly different changes in stimulus-evoked firing on hit versus miss trials (p = 0.40, p = 0.52, Wilcoxon signed-rank test, respectively). Similarly, neither RS nor FS units in striatum exhibited significantly diference changes in stimulus-evoked firing on hit versus miss trials (p = 0.12, p = 0.80, Wilcoxon signed-rank test, respectively). Finally, in amygdala, FS units showed significantly greater stimulus-evoked increases in firing rate on hit trials (7.1 ± 1.3 Hz) than on miss trials (4.0 ± 0.9 Hz; p = 5.9 x 10^-3^, Wilcoxon signed-rank test), while there was no significant difference for RS units (15.4 ± 5.6 Hz for hit, 10.0 ± 3.2 Hz for miss; p = 0.33, Wilcoxon signed-rank test). Many neurons also exhibited reward-related changes in firing rate. On hit trials, some neuronal populations exhibit rapid increases in firing rate immediately following reward (vertical dashed lines), with the most prominent increases in firing rate occurring in FS units in striatum, amygdala, and PFC. These differences in firing rates among regions, trial outcomes, and cell types were observed following stimulus presentation, but firing rates during baseline activity alone in any given cell population were not predictive of trial outcome.

In addition to single-unit activity, we recorded LFPs from all channels across the probes (**Fig. 2F-H**). Looking at averaged spectra, there was little difference in the spectra of baseline LFPs (1 s prior to stimulus) between the different trial outcomes (**Fig. 2F**). Averaging spectra like this can mask features like local peaks. For instance, recordings from PFC during 2 sessions had a low magnitude slope (**Fig. 2G**) and masked local peaks in the PFC spectra from other sessions. We therefore employed a method which separately parameterized the aperiodic components (1/f-like dynamics) and periodic components (local peaks) of LFP spectra (Donoghue et al., 2020), extracting aperiodic and periodic components in baseline LFP spectra averaged across trials of each outcome in each session (**Fig. 2G**). Periodic components in the high theta - low alpha range (center frequencies ∼7.5 Hz) were most common during baseline activity across sessions and regions; however, on average the center-frequency of local peaks in the spectra tended to be higher (high alpha or beta frequencies) prior to miss trials compared to other outcomes, particularly in PFC (p = 0.058 in S1, p = 0.032 in PFC, p > 0.2 elsewhere, one-way ANOVA). We also observed significant differences in in the slope and offset of the aperiodic components and relative power and bandwidth of the periodic components between regions (**Fig. 2H).** We computed current source density in S1 across go stimulus trials to validate and adjust where necessary our assignment of cortical layers (**Fig. 2I**). Since we were particularly interested in the relationship between spiking activity and the phase of alpha oscillations, we measured the average phase shift in the alpha band between a reference electrode (in layer 4 for the S1-striatum-amygdala implants and in prelimbic cortex (PL) for the PFC implants) and all other electrodes along the probe during baseline activity (3 s prior to stimulus) across all trials (**Fig. 2J**). There was little phase shift in the alpha band within S1 (except on the most superficial electrodes) and PFC relative to the reference electrodes, but there was a fairly uniform phase shift of ∼1 radian in striatum and amygdala relative to their reference electrode in layer 5 of S1.

### Alpha phase modulation of single-unit spiking across brain regions

Phase modulation of single neuron spiking activity varied in prevalence by brain region. Alpha modulation was commonplace in neocortex and comparatively rare in subcortical regions (**Fig. 3B**). Phase modulated units were more common in S1 (67.8 ±3.0% per session) than in striatum (13.7 ± 2.6%) or amygdala (26.2 ± 8.1%; p = 6.0 x 10^-5^, p = 5.4 x 10^-4^. Wilcoxon signed-rank test, respectively). Phase modulated units were also more common in PFC (43.6 ± 2.6% per session) than in striatum or amygdala (p = 1.3 x 10^-5^, p = 0.029, Mann-Whitney U-test, respectively). Phase modulated units were significantly more prevalent in S1 than in PFC (p = 2.1 x 10^-5^, Mann-Whitney U-test). There were also differences in the prevalence of phase modulation between RS and FS units within and between regions. Only in PFC was the prevalence of phase modulation in FS units (77.1 ± 6.5% per session) significantly higher than in RS units (40.0 ± 2.6%). In other regions, there were no significant differences between RS and FS units in the prevalence of phase modulation.

**Figure 3:**
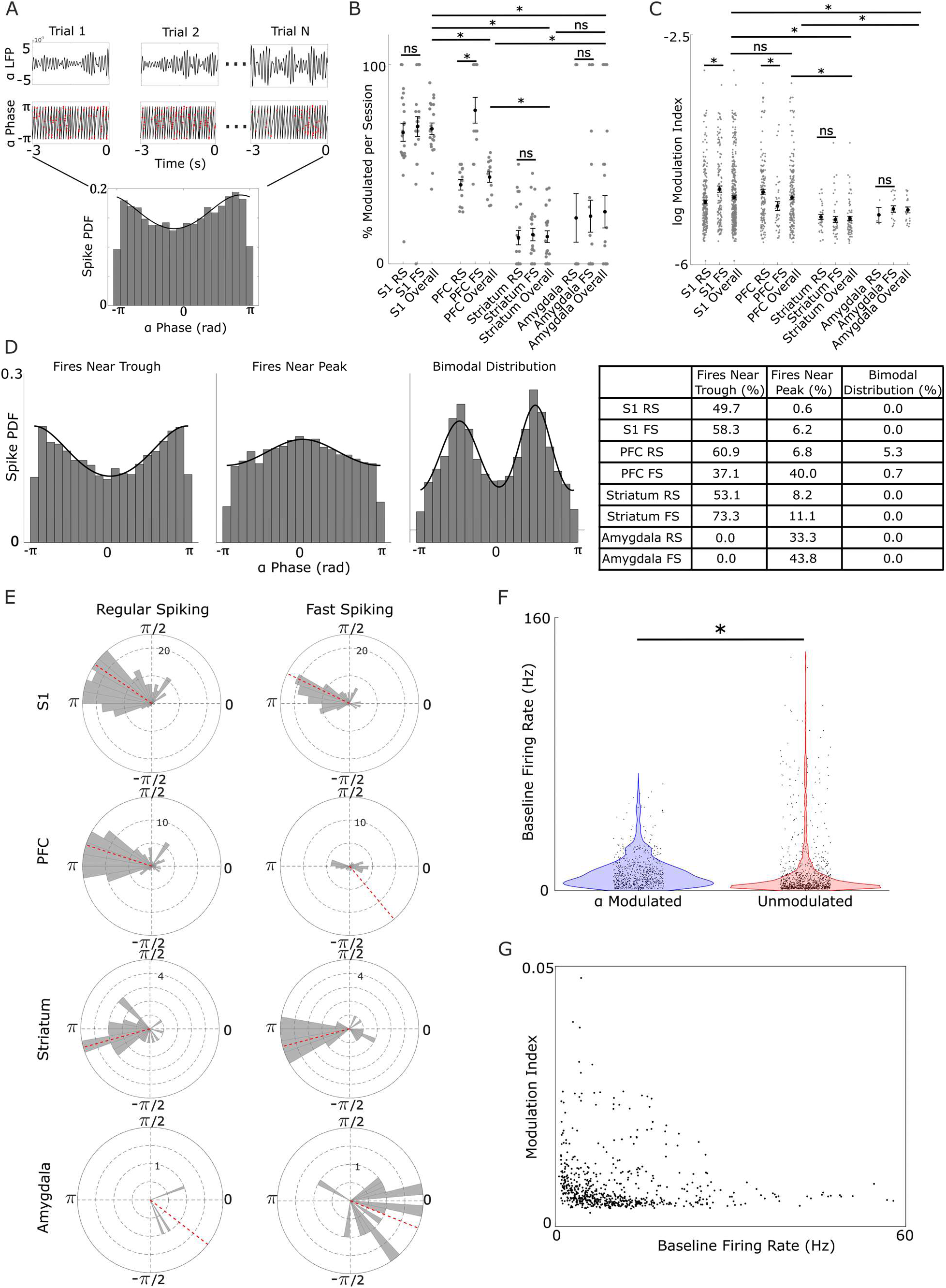
Spontaneous alpha phase modulation of single-unit activity varies across brain regions and cell types. **(A)** Examples of phase modulation of spiking activity during baseline activity on three trials. (Top) 3 s alpha band (7-12 Hz) filtered LFP. (Center) Instantaneous phase of filtered LFP computed via Hilbert transform (black). Spike times from a single neuron recorded on the same electrode are overlaid in red. (Bottom) Spiking probability density at binned phases of alpha across all trials during baseline activity. The best fit von Mises distribution is overlaid in black. **(B)** Percentages of recorded neurons which were significantly phase modulated per session separated by region (pooled across cell types) and cell type within region (RS or FS). **(C)** MI of significantly phase modulated neurons separated by region (pooled across cell types) and cell type within region (RS or FS). **(D)** Examples of 3 distinct spike-phase distributions: firing most often near the trough of alpha (left), firing most often near the peak of alpha (center), and bimodal (left). Table contains percentages of each class of spike-phase distribution for each cel type per region. **(E)** Preferred alpha phase for spiking for significantly phase modulated RS (left) and FS (right) neurons across regions. **(F)** Average baseline firing rate for phase modulated neurons compared to unmodulated neurons. **(G)** Modulation index has a weak negative correlation with firing rate across phase modulated neurons. * indicates p < 0.05.

The strength of phase modulation also varied across brain regions, with stronger coupling strength observed in neocortex than in subcortical regions (**Fig. 3C**). Across all significantly phase modulated neurons in S1 and PFC, there was no significant difference in MI (p = 0.30, Mann-Whitney U-test, respectively). However, MI in phase modulated neurons across S1 and PFC was significantly greater than MI in striatal phase modulated neurons (p = 3.5 x 10^-10^, p = 6.2 x 10^-10^, Mann-Whitney U-test, respectively). There was also a trend toward lower MI in amygdala compared to cortical regions, but this was not significant, likely due to the low number of significantly phase modulated neurons identified in amygdala. In neocortex, there were also significant differences in modulation strength between RS and FS units. In S1, MI was significantly higher in FS units than RS units (p = 0.008, Mann-Whitney U-test), while in PFC, MI was significantly higher in RS units than in FS units (p = 0.008, Mann-Whitney U-test).

Phase modulated neurons exhibited a variety of distributions of spikes relative to alpha phase (spike-phase distributions, **Fig. 3D**). The overwhelming majority of neurons had spike-phase distributions that were well characterized by the von Mises distribution (analogous to a normal distribution in circular space). Most of these neurons (49.2% of significantly phase modulated neurons) spiked most often near the trough of the alpha oscillation, consistent with previous findings of alpha modulation of cortical multi-unit spiking in nonhuman primates (Bollimunta et al., 2008; Haegens et al., 2011; van Kerkoerle et al., 2014; Dougherty et al., 2017). Some of these neurons (9.1% of significantly phase modulated neurons) spiked most often near the peak of the alpha oscillation. In neocortex, all neurons that spiked near the peak of the alpha oscillation were located in infragranular layers. Most of the remaining neurons fired most often at intermediate phases of the alpha oscillation but were still well characterized by the von Mises distribution. However, a handful of neurons (2.2% of phase modulated neurons), all located in PFC, exhibited bimodal spike-phase distributions which were better characterized by a mixture model of two von Mises distributions. In neocortex, we observed significant differences in the distributions of preferred phase between RS and FS neurons (p = 0.02 in S1, p = 0.001 in PFC, Kuiper’s test) (**Fig. 3E**). Conversely, there were no significant differences in preferred phase distributions between RS and FS neurons in subcortical regions.

The distributions of baseline firing rates between significantly alpha modulated neurons and unmodulated neurons were largely overlapping but did differ significantly (p = 1.2 x 10^-19^, Mann-Whitney U-test, **Fig. 3F**). Although the mean baseline firing rates of phase modulated and unmodulated neurons were comparable, the variance was much higher in unmodulated neurons (1.1 ± 3.8 Hz, 1.1 ± 6.0 Hz, respectively). Specifically, neurons with very high baseline firing rates (>60 Hz) were less likely to be phase modulated.

Compared to other regions, we recorded very few significantly phase modulated RS or FS units in amygdala (3 and 9 in total across all sessions, respectively). Due to the low number of units recorded in amygdala (74 across all sessions), and the paucity of significantly phase modulated units among them, we have restricted further analyses to neocortical and striatal neurons.

#### Phase modulated neurons exhibit enhanced responsiveness to task-relevant stimuli

Phase modulated neurons tended to exhibit greater stimulus-induced and task-related changes in spiking activity compared to their unmodulated counterparts. This was observed in the average change in firing rate from baseline firing rate (3s prior to stimulus onset) across different populations, especially in FS units (**Fig. 4A**). To quantify stimulus-evoked changes in firing rate, we limited our analysis to differences in average firing rate during baseline and average firing rate during stimulus presentation (0 – 0.2 s) **(Fig. 4B).** At the population level, phase modulated FS units in neocortex and RS units in striatum responded more strongly to contralateral tactile stimuli than their unmodulated counterparts. In S1 and PFC, phase modulated FS units exhibited greater stimulus-evoked changes in firing rate compared to unmodulated FS units on hit trials (p = 2.7 x 10^-5^, p = 3.2 x 10^-5^, Mann-Whitney U-test, respectively). RS units in S1 and PFC, however, did not exhibit significant differences in stimulus-evoked changes in firing rate on hit trials. Conversely, phase modulated RS, not FS, units in striatum exhibited significantly greater stimulus-evoked changes in firing rate on hit trials compared to their unmodulated counterparts (p = 0.002, Mann-Whitney U-test).

**Figure 4:**
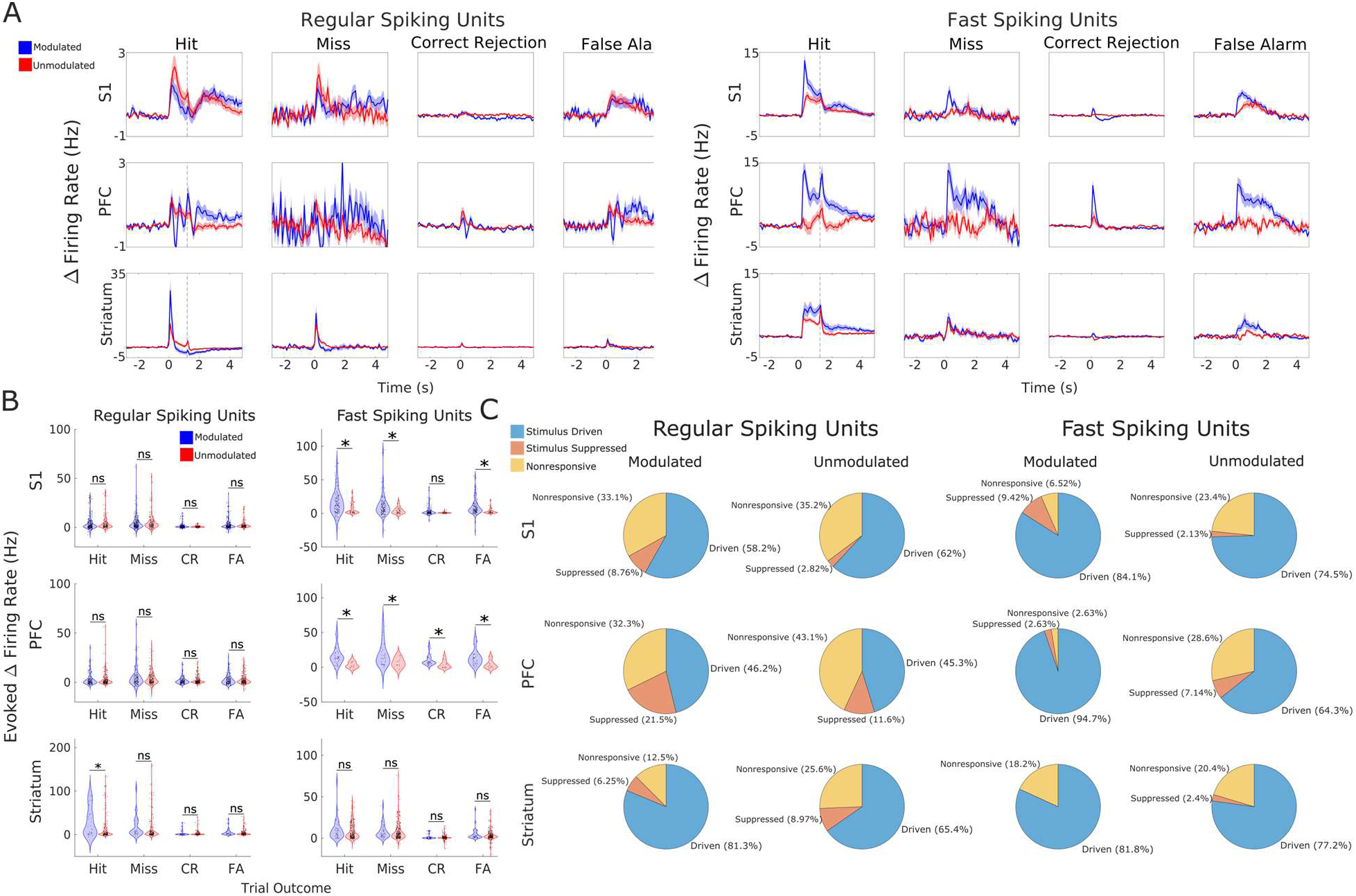
Phase modulated neurons produced stronger responses to tactile stimuli than unmodulated neurons. **(A)** Change in firing rate from baseline (calculated across 3 s prior to stimulus presentation) averaged across populations for each trial outcome for alpha modulated (blue) and unmodulated (red) neurons. **(B)** Evoked change in firing from baseline by tactile stimulus during 0.2 s period following stimulus onset for each region, population, and trial outcome. **(C)** Pie charts representing percentage of modulated and unmodulated neurons that were driven (blue), suppressed (orange), or unresponsive to contralateral tactile stimuli on hit trials. * indicates p < 0.05.

Interestingly, phase modulated FS neurons in neocortex also tended to exhibit greater changes in firing rate in response to ipsilateral tactile stimuli compared to unmodulated FS neurons (**Fig. 4B**). In S1 phase modulated FS neurons, there was a trend toward greater stimulus-evoked changes in firing rate on correct rejection trials (p = 0.051, Mann-Whitney U-test), and in PFC phase modulated FS neurons there was significantly higher stimulus-evoked changes in firing rate compared to unmodulated FS neurons on correct rejection trials (p = 0.002, Mann-Whitney U-test) FS units in S1 and PFC exhibited greater stimulus-evoked changes in firing rate than unmodulated FS units on miss and false alarm trials as well (p < 0.001, Mann-Whitney U-test).

Not all neurons respond to go stimuli with increased firing rate; some neurons showed suppressed firing rates in response to contralateral tactile stimulation, and others did not alter their firing rates at all. We therefore classified neurons as driven (average firing rate increase of > 1 Hz from baseline within 0.2 s of stimulus across hit trials), suppressed (average firing rate decrease of > 1 Hz from baseline within 0.2 s of stimulus across hit trials), and nonresponsive (remaining neurons, **Fig. 4C**). Phase modulated FS units in S1 and PFC and phase modulated RS units in PFC were significantly more likely to be responsive (driven or suppressed) than their unmodulated counterparts (p < 0.05, Fisher’s exact test).

#### Phase modulation varies across cortical layers and subregions

The prevalence and strength of phase modulation varied across cortical layers in S1 (**Fig. 4A**). Approximate layers in S1 were assigned by monitoring electrode position in real-time using Neuropixels Trajectory Explorer during implantation (see Methods). We then validated those layer assignments and adjusted where appropriate using current source density (CSD) estimation from stimulus evoked changes in LFP activity (**Fig. 2H**). The fraction of both RS and FS units which were modulated by alpha phase per session differed significantly across cortical layers (p = 0.005, p = 4.5 x 10^-4^, one-way ANOVA, respectively) (**Fig. 4A**). Phase modulated RS units were most common in layer 4. Phase modulated FS units, by contrast, were most common in layers 1-3 and became progressively less common in more ventral layers. The high incidence of alpha phase modulation of RS units in granular cortex compared to infragranular cortex is consistent with previous observations in macaque primary visual cortex (Dougherty et al., 2017). MI varied significantly across cortical layers for both RS (p = 7.9 x 10^-5^, one-way ANOVA) and FS units (p = 1.7 x 10^-9^, one-way ANOVA), with the strongest alpha modulation observed in layer 1 FS units (0.018 ± 0.001). Finally, the preferred alpha phase of spiking changed significantly with cortical depth (RS: p = 1.8 x 10^-7^, FS: p = 4.9 x 10^-9^, Watson-Williams test). As previously noted, only neurons in infragranular layers fired near the peak of the alpha oscillation, and these drive the average preferred phase in these layers toward 0 radians. We hypothesize that the FS neurons whose preferred phase was near 0 radians are the source of rhythmic inhibition that generates the more common variety of phase modulation in neocortex.

Probes implanted in PFC spanned a number of subregions in PFC but were located entirely in infragranular layers. We therefore could not use CSD to validate our subregion assignments in PFC as we did for layers in S1, so our subregion assignments are putative. The prevalence of phase modulation also varied significantly between subregions of PFC among RS (p = 4.2 x 10^-4^, one-way ANOVA) and FS units (p = 0.005, one-way ANOVA) (**Fig. 4B**). Phase modulated RS units were rare in anterior cingulate cortex (ACC; 7.6 ± 4.7% across sessions) and orbitomedial cortex (ORB; 6.7 ± 6.7%) and most common in infralimbic cortex (IL; 56.8 ± 10.4%) and dorsal peduncular area (DP; 58.8 ± 6.6%). Phase modulated FS units followed their own distribution, with high prevalence in ACC (88.1 ± 7.9%) and IL (100% across all sessions). MI varied significantly across subregions of PFC for RS and FS units (p = 1.6 x 10^-3^, p = 0.013, one-way ANOVA, respectively). Compared to laminar differences in S1, the phase of alpha at which neurons fired most often varied considerably across PFC subregions (RS: p = 2.4 x ± 10^-12^, FS: p = 2.2 x ± 10^-16^, Watson-Williams test). For RS units in ORB, FS units in DP, and units of both classes in ACC, the average preferred phase was near 0 radians (spikes occurred most frequently around the peak of the alpha oscillation).

#### Dynamic changes in alpha power and phase modulation

Since alpha power changes dynamically, even over the course of the 3 s baseline periods of activity on which our analyses have focused, we naturally wondered whether phase modulation of single-unit activity varied with alpha power (**Fig. 5**). To this end, we identified low and high alpha power events during baseline activity on each electrode where we recorded a phase modulated neuron. Low alpha power events were defined as epochs ≥ 0.3 s where alpha power remained below its 50th percentile (on that electrode across the recording session). Similarly, high alpha power events were defined as epochs ≥ 0.3 s where alpha power remained above its 75th percentile. In an example neuron, we observed different spike-phase distributions during low- and high-power events (**Fig. 5A**). Both were well characterized by a von Mises distribution, but the distribution during high alpha power events deviated much more strongly from a uniform distribution. We compared MI, average firing rate, and preferred firing phase of alpha across spikes for all phase modulated RS and FS units during low and high alpha power events (**Fig. 5B**). We also compared the percentage of units in each session which were phase modulated during low and high alpha power events. For RS and FS units across S1, PFC, and striatum, MI increased significantly between low and high alpha power events (p < 1.1 x 10^-6^, Wilcoxon signed-rank test). Interestingly, there was little difference in firing rates between low and high alpha events. Although FS neurons in S1 and RS neurons PFC exhibited significantly different firing rates between high and low alpha events (p = 7.9 x 10^-4^, p = 0.029, Wilcoxon signed-rank test, respectively), the differences between firing rates during high and low alpha events across populations are mixed in their direction (some higher during high alpha, some higher during low alpha) and the average difference in firing rate is at most 1.0 Hz (S1 FS neurons). This implies that increased alpha power does not simply reflect an increase in rhythmic inhibitory tone, which one would expect to reduce overall firing rate. Instead, the higher MI during high alpha power events, in conjunction with the modest changes in firing rates, suggest that increased alpha power reflects balanced increases in rhythmic excitation and inhibition. Finally, there was little difference in the preferred alpha phase of spiking between low and high alpha power events.

**Figure 5:**
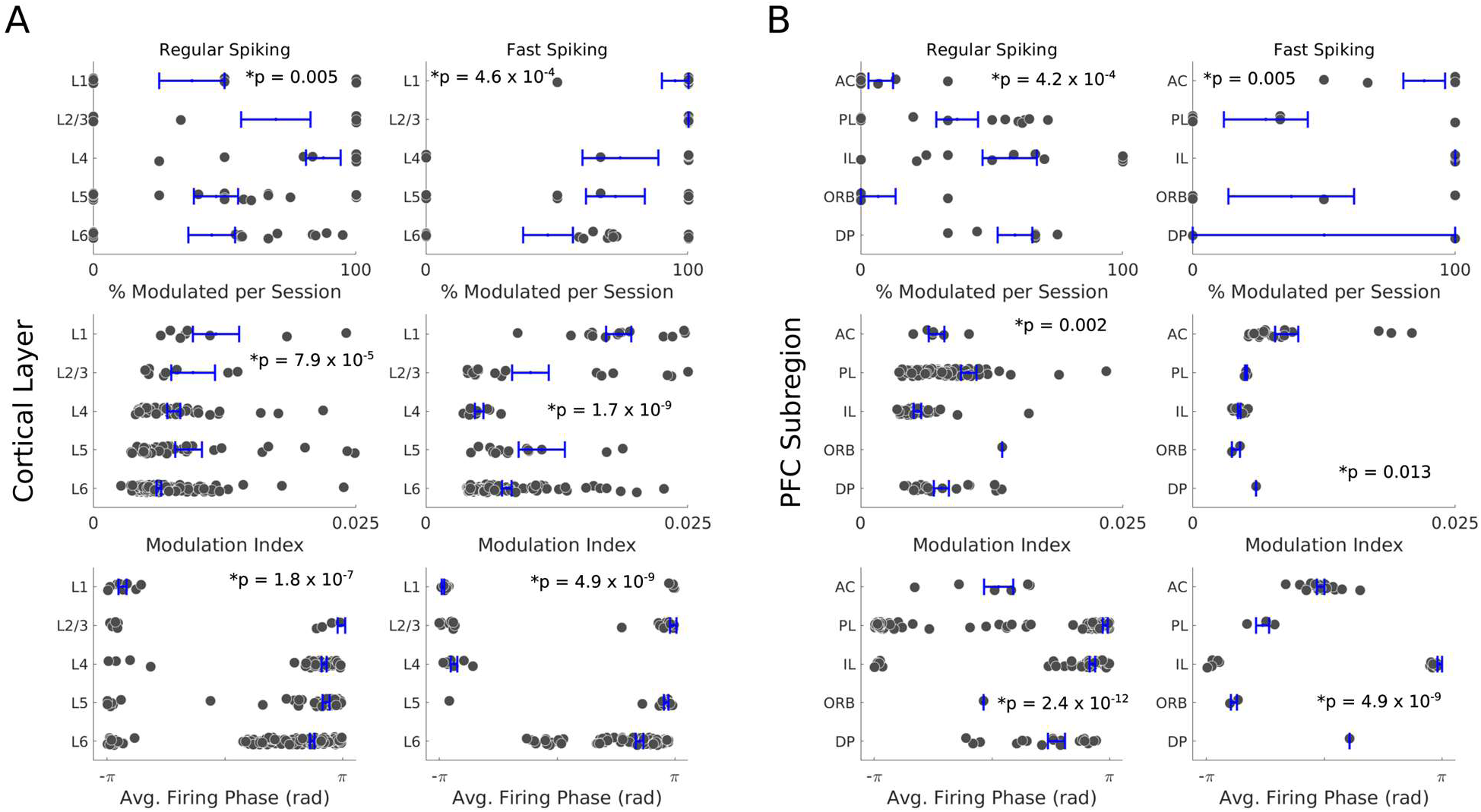
Phase modulation varies across cortical layers in S1 and subregions in PFC. (Top to bottom) Percent of RS (left) and FS (right) units that were phase modulated per session, MI for all significantly phase modulated neurons, and preferred firing phase of phase modulated neurons in different (**A**) S1 cortical layers and (**B**) PFC subregions. * indicates p < 0.05.

#### Single neuron phase modulation and its relation to behavior

We next investigated how spontaneous (pre-stimulus) alpha modulation of single neuron spiking activity related to cognitive performance (correct vs. incorrect) and motor activity (response vs. no response) during the selective detection task. To quantify these relationships, for each neuron previously identified as alpha modulated, we pooled the instantaneous alpha phase of each spike across trials of a given outcome: hit, miss, correct rejection, false alarm, correct (hit and correct rejection), incorrect (miss and false alarm), response (hit and false alarm), and no-response (miss and correct rejection). We then computed p-values from the distributions of baseline spike phases of each trial outcome using Rayleigh’s test and used the same corrected p-value threshold as above to determine if the neuron’s spiking activity was significantly modulated by alpha phase prior to trials of a given outcome. **Figure 7A** shows an example of an alpha modulated neuron and its spike-phase distributions for different cognitive outcomes (correct and incorrect) and motor actions (response and no-response). This example neuron exhibited a common motif: significant alpha modulation prior to correct trials, response trials, and no-response trials, but not prior to incorrect trials.

**Figure 6:**
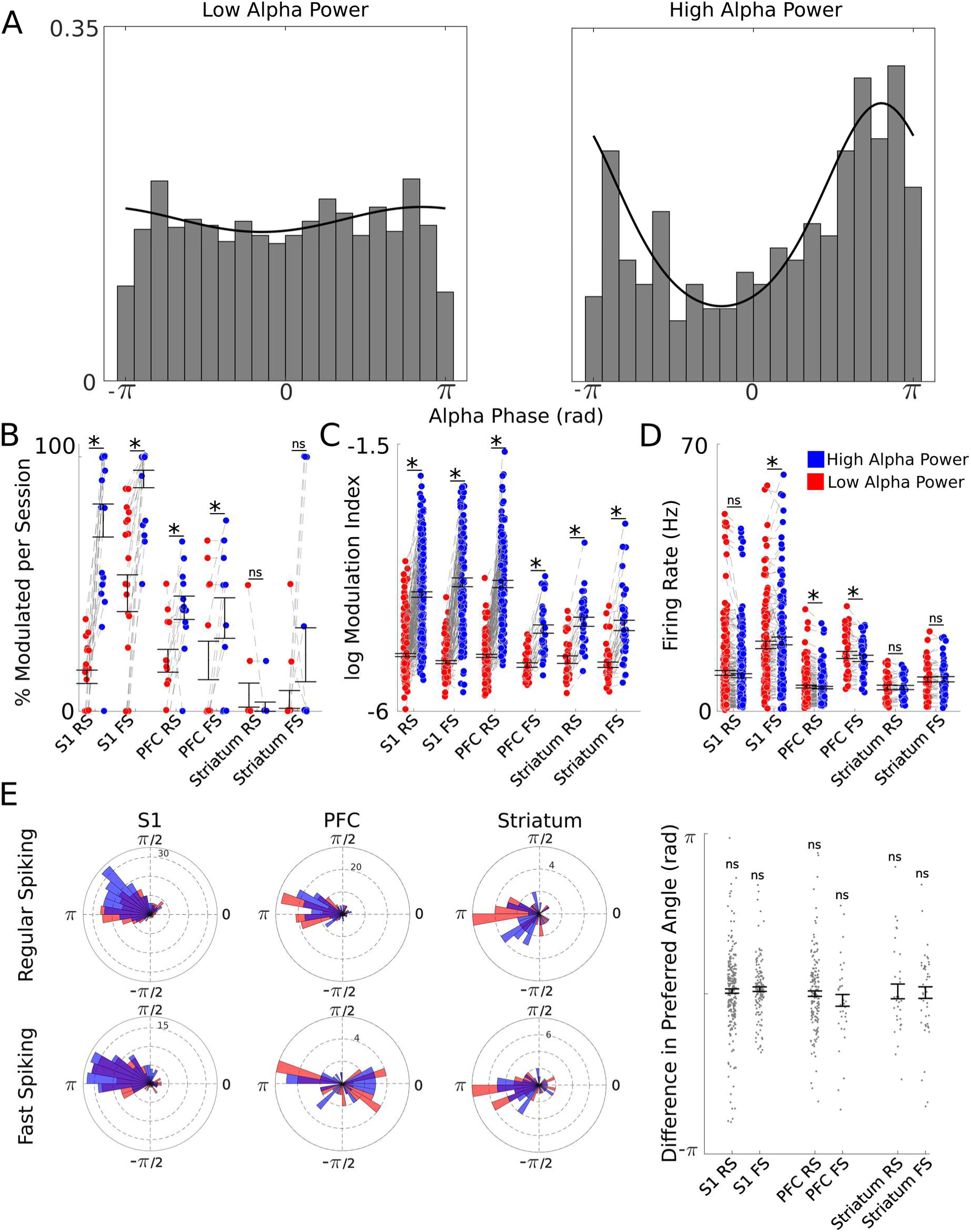
Phase modulation of single-unit activity changes with alpha power. **(A)** Example of spike-phase distributions across alpha phase during low alpha power events (left, ≥ 0.3 s when alpha power is below its 50th percentile) and high alpha power events (right, 0.3 s when alpha power is above its 75th percentile) for a single neuron. **(B)** Percent of phase modulated neurons (determined across all trials) that were significantly phase modulated across low alpha power events (red) and high alpha power events per session. **(C)** Average MI and **(D)** average firing rate of phase modulated neurons during low and high alpha power events. **(E)** Preferred alpha phase of phase modulated neurons during low and high power alpha events (left) and the difference in preferred phase angles between high and low alpha power events (right). * indicates p < 0.05.

**Figure 7:**
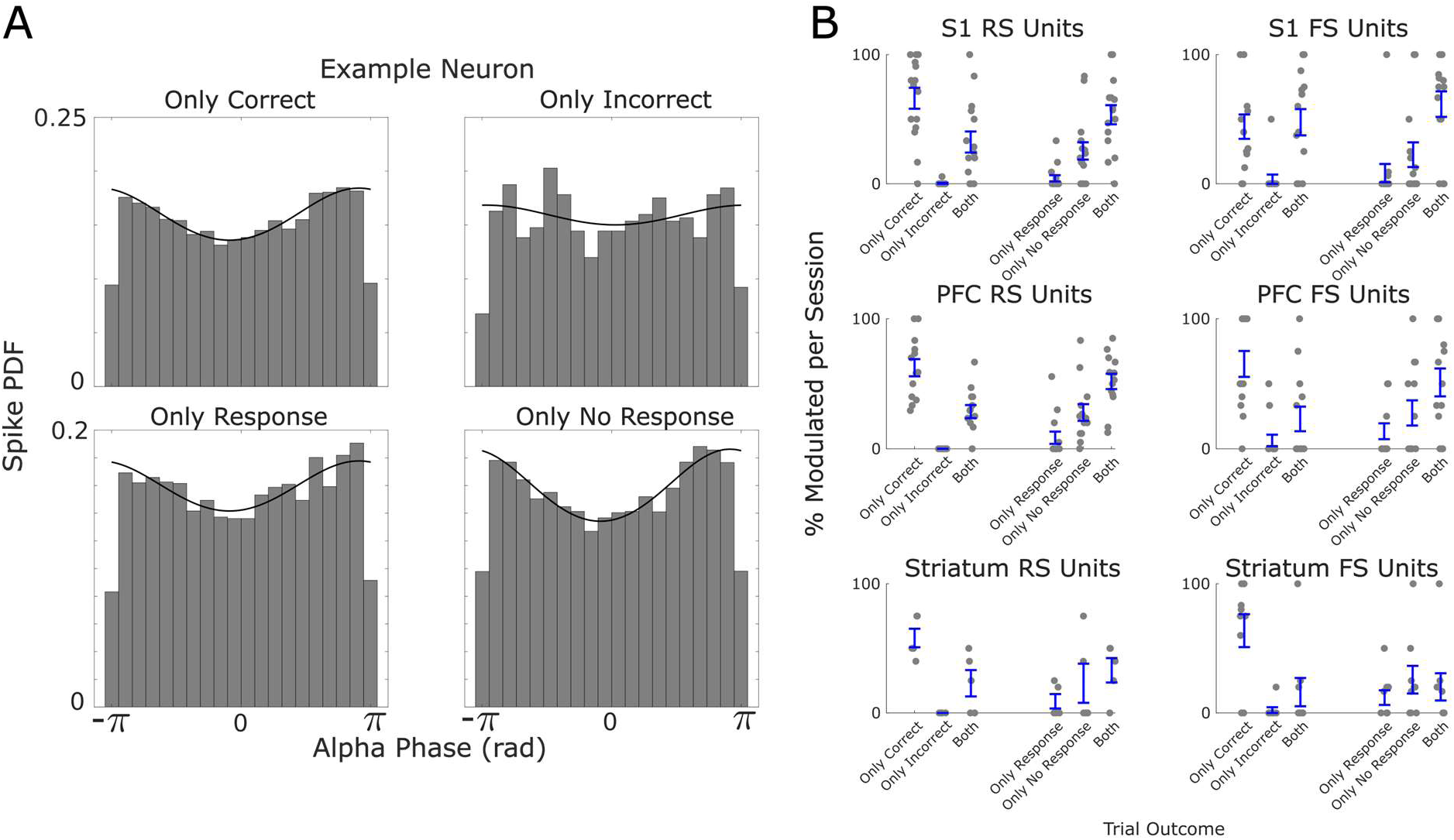
Incorrect trials were preceded by a disruption in alpha phase modulation. **(A)** Spike-phase distributions for baseline activity prior to correct (hit and correct rejection), incorrect (miss and false alarm), response (hit and false alarm), and no-response (miss and correct rejection) trials for an example neuron. Only prior to incorrect trials was this neuron not significantly phase modulated. **(B)** Percent of phase modulated neurons which were significantly phase modulated prior to trials of a given outcome. (Left) Percent of phase modulated neurons (determined across all trials) which were either only phase modulated prior to correct trials, only phase modulated prior to incorrect trials, or significantly phase modulated prior to both correct and incorrect trials. (Right) Percent of phase modulated neurons (determined across all trials) which were either only phase modulated prior to response trials, only phase modulated prior to no-response trials, or phase modulated prior to both response and no-response trials.

To systematically investigate the relationship between alpha phase modulation and cognitive performance, we computed the proportion of phase modulated units of each waveform class (RS or FS) which were selectively phase modulated on correct trials, selectively phase modulated on incorrect trials, or phase modulated regardless of the cognitive outcome of the trials (i.e., phase modulated prior to both correct and incorrect trials) per session. Similarly, to investigate the relationship between alpha phase modulation and motor output, we computed the proportion of phase modulated units of each waveform class that were selectively phase modulated on response trials, selectively phase modulated on no-response trials, or phase modulated regardless of motor outcome (i.e., phase modulated prior to both response and no-response trials) (**Fig. 7B**). Neurons which were selectively phase modulated prior to correct trials were the most common across brain regions, while neurons which were phase modulated regardless of trial outcome were less common, and neurons which were selectively phase modulated prior to incorrect trials were extremely rare. By comparison, the differences in proportions of neurons selectively phase modulated prior to response or no-response trials, or those that were phase modulated prior to both, were less pronounced than when focusing on cognitive outcome.

Since the mice included in this study were experts at the selective detection task, there were more correct trials included in our analyses than incorrect trials. To verify that the high incidence of neurons selectively phase modulated prior to correct trials was due to a reduction of phase modulation prior in incorrect trials, rather than an artifact of the smaller number of incorrect trials and Rayleigh’s test being therefore underpowered, we employed a bootstrapping method (**Supp.** Fig. 1A). For each neuron which was selectively phase modulated prior to correct, but not incorrect, trials, we performed Rayleigh’s test on a random selection of correct trials where the number correct trials included was matched to the number of incorrect trials in that session. We repeated this process 100 times with different random selections of correct trials to generate a distribution of p-values. If the Rayleigh’s test p-value associated with incorrect trials was greater than the 95^th^ percentile of the trial-matched correct trial p-value distribution, we considered that neuron to exhibit significantly reduced phase modulation prior to incorrect trials compared to correct trials. Of the neurons selectively phase modulated prior to correct trials, 18.4% exhibited a significant reduction in phase modulation prior to incorrect trials (**Supp.** Fig. 1B). By contrast, only 1.4% had a Rayleigh’s p-value across incorrect trials less than the 5^th^ percentile of the trial-matched correct trial p-value distribution. Taken together, these data suggest that poor cognitive performance on the task was preceded by disruptions in the modulation of single neuron activity by the phase of alpha oscillations.

Finally, we investigated whether spontaneous motor activity, specifically spontaneous licking during the inter-trial period, impacted alpha modulation of single-unit spiking activity (**Fig. 8**). Inter-trial intervals containing spontaneous licking were significantly less common than those without spontaneous licks (p = 3.7 x 10^-7^, Wilcoxon signed-rank; **Fig. 8A**). In line with previous work, spontaneous motor activity correlated with reduced task performance (Marrero et al., 2022). Here, we observed significantly reduced discriminability on trials preceded by spontaneous licking compared to those without (p = 0.015, Wilcoxon signed-rank; **Fig. 8B**). Most phase modulated neurons (determined across all trials) were significantly phase modulated across only trials without spontaneous licks but were not significantly phase modulated across only trials with spontaneous licks (p = 3.6 x 10^-7^, Wilcoxon signed-rank; **Fig. 8C**). Because there were so many more inter-trial intervals without licks than with licks, we employed a similar bootstrapping procedure as for correct and incorrect trials (**Supp. Fig. 2**). For each phase modulated neuron, we performed Rayleigh’s test on a random selection of trials without spontaneous licks where the number trials included was matched to the number of trials with spontaneous licks in that session. 29.3% of phase modulated neurons had a Rayleigh’s p-value across spontaneous lick trials greater than the 95^th^ percentile of the trial-matched no-lick p-value distribution (significant reduction in phase modulation associated with spontaneous licking), while 15.0% of phase modulated neurons had a Rayleigh’s p-value less than the 5^th^ percentile of the trial-matched no-lick p-value distribution (significant increase in phase modulation associated with spontaneous licking) (**Supp. Fig. 2**). Interestingly, all phase modulated neuronal population exhibited significant increases in baseline firing rate on trials with spontaneous licks (p < 0.05, Wilcoxon signed-rank) with the exception of RS neurons in striatum, but none of the unmodulated populations exhibited significant increases in firing rate associated with spontaneous licking (**Fig. 8D**).

**Figure 8:**
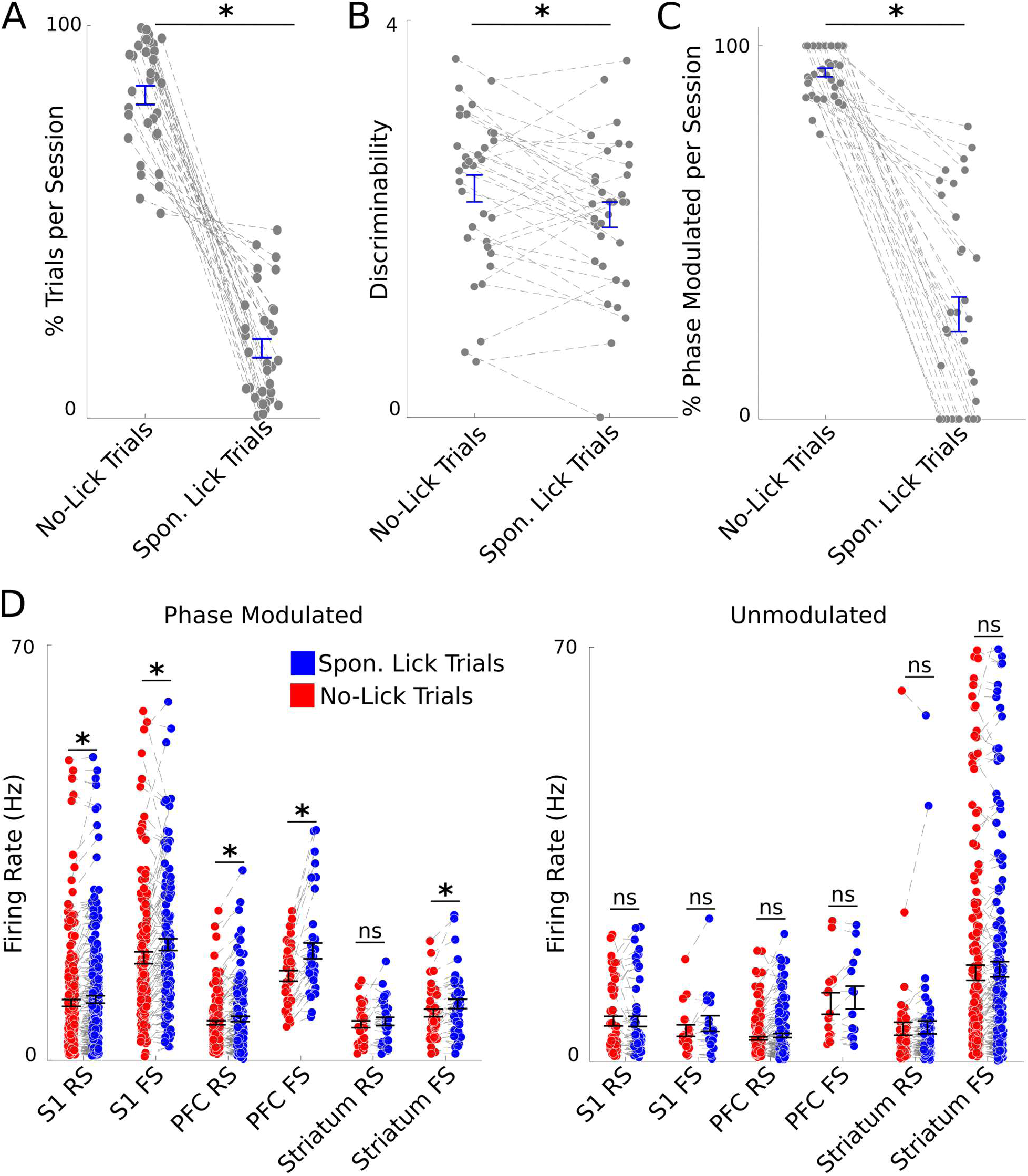
Spontaneous licking interrupts phase modulation, and phase modulated neurons exhibit greater changes in firing rate associated with spontaneous licking than unmodulated neurons. (A) Number of trials with and without spontaneous licks in each session. (B) Discriminability across trials with and without spontaneous licks in each session. (C) Percent of phase modulated neurons (determined across all trials) that were phase modulated across trials with and without spontaneous licks. (D) Baseline firing rate on trials with (blue) and without (red) spontaneous licks for all phase modulated neurons (left) and all unmodulated neurons (right). * indicates p < 0.05.

## Discussion

### Anatomy and physiology of spontaneous alpha modulation

In this study, we systematically characterized alpha modulation of single-unit spiking activity in two cortical (S1 and PFC) and two subcortical (striatum and amygdala) brain regions during baseline activity associated with different behavioral outcomes. We found phase modulated neurons in each brain region, but they were far more prevalent in neocortex than in striatum or amygdala (**Fig. 3B**). Phase modulated neurons were also more common in S1 than in PFC; however, all neurons recorded from PFC were located in infragranular layers, whereas neurons recorded from S1 were distributed across all cortical layers. Within PFC, we found significant differences in the prevalence of phase modulation among its subregions (ACC, PL, IL, etc.; **Fig 5B**). Differences in the prevalence of alpha modulation between cortical regions have also been reported in macaques (Bollimunta et al., 2008; Haegens et al., 2011).

Within S1, we found the prevalence and strength of alpha modulation differed significantly between cortical layers (**Fig. 5A**). Studies of phase modulation of evoked multi-unit spiking in nonhuman primates also observed laminar differences in the prevalence of phase modulation (Bollimunta et al., 2008; Dougherty et al., 2017; Davis et al., 2023). Bollimunta et al. (2008) observed the weakest coupling between alpha and multi-unit activity in supragranular layers of higher order visual areas (V2 and V4) in macaques, while Davis et al. (2023) observed the strongest coupling between generalized oscillatory phase (5 - 50 Hz) and multi-unit spiking across PFC and medial temporal lobe in marmosets and V4 of macaques. We observed very sparse alpha modulation among RS units in layer 1, but most neurons in layer 2/3 were alpha modulated. It may be that multi-unit activity recordings in supragranular cortex from Bollimunta et al. (2008) were dominated by layer 1 neurons, and recordings from Davis et al. (2023) were dominated by layer 2/3 neurons, if in fact the laminar distributions of phase modulation are comparable between mice and nonhuman primates.

The majority of phase modulated neurons exhibited spike-phase distributions which closely followed a von Mises distribution (a normal distribution in circular space). In other words, these neurons preferentially fired at one particular phase of the alpha cycle (**Fig 3D**). In neocortex, most neurons fired most often at the trough of the alpha cycle (**Fig. 3D, E**) in line with previous findings of alpha modulation of cortical multi-unit spiking in nonhuman primates (Bollimunta et al., 2008; Haegens et al., 2011; van Kerkoerle et al., 2014; Dougherty et al., 2017). This was also true for phase modulated neurons in striatum (**Fig. 3D, E**). In S1 and PFC, a small fraction of both RS and FS units preferentially fired near the peak of the alpha cycle, however (**Fig. 3D, E**). All of the neurons in neocortex that preferentially fired near the peak of the alpha cycle were located in infragranular layers. Neocortical FS neurons that preferentially fired near the peak of the alpha cycle were likely GABAergic interneurons which generated the pulsed inhibition necessary for the more common variety of alpha modulation (Mazaheri and Jensen, 2010). The RS units that preferentially fired at the peak of alpha may have been somatostatin-expressing interneurons, which are commonly found in layer 5 (Xu et al., 2010; Riedemann, 2019), also generating pulsed inhibition. Alternatively, those RS units may have been layer 5 pyramidal neurons expressing h- and T-currents which can both generate and pace the alpha rhythm (Jones et al., 2000). There were a number RS and FS units in striatum that preferentially fired near the peak of the alpha cycle (**Fig. 3E**), but it is unclear whether these neurons generated alpha modulation locally or if their modulated spiking distributions were inherited from elsewhere, particularly neocortex. The small number of phase modulated neurons in amygdala often preferentially fired at neither the peak nor the trough of the alpha oscillation. Finally, a small number of neurons in PFC exhibited a bimodal distribution with respect to alpha phase (**Fig. 3C**). Decreased firing near the peak of the alpha oscillation in these neurons is consistent with the pulsed inhibition hypothesis. The symmetrical intermediated phases of alpha at which they preferentially spiked suggests the are driven by excitatory connections from neurons that are themselves coupled to the alpha rhythm. The mechanism by which this form of coupling arises remains an open question.

Phase modulation of baseline spiking activity correlated strongly with tactile stimulus encoding. Phase modulated neurons, particularly FS units, across brain regions exhibited greater task-related changes in firing rate from baseline than their unmodulated counterparts (**Fig. 4A, B**). We hypothesize that the pulsed inhibition responsible for alpha modulation kept stimulus-driven neurons’ firing rates near the minimum of their dynamic ranges such that tactile stimuli produced greater changes in firing rate. In this sense, the pulsed inhibition associated with alpha oscillations recruits a population of neurons primed to encode sensory stimuli, consistent with the alpha rhythm’s association with a prepared brain state (Adrian and Matthews, 1934; Nikouline et al., 2000; Fanselow et al., 2001; Cooper et al., 2003; Tort et al., 2010b). The finding that unmodulated neurons were more likely to be unresponsive to contralateral tactile stimuli (**Fig. 4C**) is consistent with previous work showing a subpopulation of FS units in S1 which were unresponsive to tactile stimuli and fired steadily at gamma frequencies (Shin and Moore, 2019). These neurons fire with such regularity and at such high frequency they could not be modulated by alpha rhythms.

Our findings suggest that phase modulated neurons receive more diverse synaptic inputs than their unmodulated counterparts. Previous work has demonstrated sparse encoding of ipsilateral (non-aligned) tactile stimuli in S1 of awake mice mediated by corticocortical projections via the corpus callosum (Pala and Stanley, 2022). Here we show that phase modulated FS units, especially in neocortex, encode ipsilateral tactile stimuli than unmodulated FS units (**Fig. 4A, B**). We also observed that alpha modulated neurons, particularly in PFC, exhibited higher firing rates following reward delivery compared to their unmodulated counterparts **(Fig. 4A**). Whether these elevated firing rates are related to reward processing or simply the motor activity associated with licking to collect the water reward, this increased activity suggests that neurons modulated by alpha phase tend to be innervated by more diverse sources than sensory stimuli alone. Additionally, phase modulated neurons exhibited significantly higher baseline firing rates on trials with spontaneous licking prior to stimulus presentation, while unmodulated neurons did not (**Fig. 8D**). This increased firing rate associated with spontaneous motor activity provides further support for the notion that phase modulated neurons are driven by more diverse presynaptic populations.

We observed stronger alpha modulation of spiking activity during events with sustained high alpha power than during events with sustained low alpha power (**Fig. 5**), which was aligned with predictions from the pulsed inhibition hypothesis of alpha oscillations (Klimesch et al., 2007; Mazaheri and Jensen, 2010; Mathewson et al., 2011). However, the increased alpha modulation from low to high alpha events was accompanied by negligible changes in firing rate and preferred phase (**Fig 5C**). This suggests that increased pulsed inhibition during high alpha events may be accompanied by balanced, antiphase pulses of excitation, requiring an amendment to the pulsed inhibition hypothesis, at least insofar as it applies to mice.

#### Spontaneous alpha modulation and behavior

In this study, we investigated how alpha modulation during baseline activity correlated to performance of a whisker-based selective detection task. Consistent with previous results showing that pre-stimulus alpha modulates sensory encoding in human perceptual-decision making (Lou et al., 2014), we found significant deficits in alpha modulation prior to incorrect (miss or false alarm trials) trials compared to correct (hit or correct rejection) trials (**Fig. 7**). While there were neurons that were significantly phase modulated prior to both correct and incorrect trials, most neurons across regions were selectively phase modulated prior to correct trials, with extremely few neurons showing selective phase modulation prior to incorrect trials (**Fig. 7B**). Furthermore, many neurons which were selectively phase modulated prior to correct trials exhibited significant reductions in phase modulation prior to incorrect trials as evaluated through a boostrapping approach (**Supp.** Fig. 1). Thus, a dysregulation of alpha modulation of spiking activity during spontaneous activity correlated with poor performance on the selective detection task. This may reflect a lapse in attention.

Similar to many tactile tasks, the behavioral task employed in this study involves the coordinated activity of numerous brain regions, including thalamus, S1, motor cortex, anterior lateral motor cortex, and striatum (Zheng et al., 2015; Delis et al., 2018; Rodenkirch et al., 2019; Aruljothi et al., 2020; Zareian et al., 2021; Marrero et al., 2022; Zareian et al., 2023). Marrero et al. (2022) found that correct responses to sensory stimuli were predicted by a pre-stimulus global low amplitude state across cortex with reduced spontaneous activity, principally identified through widefield calcium imaging of activity in dorsal neocortex. The low activity state associated with correct task performance was most prominent in neurons that encode stimuli or were associated with motor response. This suggests that a quiet brain state is preferable for neural circuits to process information suited for detection tasks, likely at the cost of discrimination performance (Wang et al., 2010; Ollerenshaw et al., 2014). Alpha oscillations have been hypothesized to produce a brain state which is “calm yet alert” (Adrian and Matthews, 1934; Cooper et al., 2003), which is in line with strong alpha modulation associated with correct responses on the selective detection task.

Buffalo et al. (2011) evaluated the coherence between multi-unit activity and LFPs, which is comparable to phase modulation, in visual cortex of macaques performing a cued visual attention task and found reduced coherence between multi-unit activity and low frequency oscillations (including alpha) while the monkey attended to visual stimuli. At first, this may seem to contradict our observation of weaker alpha modulation prior to incorrect trials; however, in the visual attention task, nonhuman primates were cued to attend to the visual stimulus (requiring internal direction of attention), while the mice performing the tactile selective detection task received no cue prior to stimulus presentation. The tasks were therefore sufficiently distinct that too close a comparison would be inappropriate. The selective detection task employed here does not require the fine discrimination tactile stimuli; instead, it requires all-or-none responses to stimuli distributed across much of the whisker pad. The strong alpha modulation observed prior to correct trials compared to incorrect trials, rather than reflecting the sort of internally directed attention described in Buffalo et al. (2011), instead may produce a prepared state which tunes brain networks to the discrete all-or-none behavior required for this task (Hô and Destexhe, 2000).

A number of studies have associated disruptions in phase modulation of spiking activity in pathological and pharmacologically altered brain states (Sigurdsson et al., 2010; Lopez-Pigozzi et al., 2016; Lazaro et al., 2019; Golden and Chadderton, 2022; Weiss et al., 2023). Golden and Chadderton (2022) injected mice with psilocybin and observed reduced coupling of single-unit spiking activity to different LFP bands (delta, theta, alpha, and beta) in ACC. While that study focused on the effects of psilocybin on the physiology of anterior cingulate rather than mouse behavior, it is interesting to note that attention deficits associated with psilocybin more likely reflect a decreased ability to suppress and ignore distraction than a reduction in attentional capacity per se (Carter et al., 2005), and alpha oscillations are strongly implicated in the suppression of distractors (Cooper et al., 2003; Klimesch et al., 2007; Palva and Palva, 2007; Lazaro et al., 2019). Lazaro et al. (2019) observed alterations in phase modulation in PFC of a mouse model of autism (CNTNAP2 KO). While they did not link altered phase modulation, or disruption of network dynamics more generally, to specific behavioral differences, it is noteworthy that autism is associated with cognitive and attentional deficits (Cantio et al., 2016; Muskens et al., 2017). Taken together, these studies suggest that alpha modulation is a feature of a properly functioning cortex and subject to pathological and pharmacological dysfunction. A recent study developed a real-time system for stimulating neural populations at particular phases of LFP oscillations (Wick et al., 2023). Future work may harness such techniques to investigate whether performance in a task like this can be improved by stimulating cortical neurons at appropriate phases of the alpha rhythm during spontaneous activity.

## Acknowledgements

This work was supported by AFOSR FA9550-22–1-0337, NSF CBET 1847315, NIH R01NS119813, and NIH R01AG075114.

## Disclaimer

Q.W. is the co-founder of Sharper Sense.

## Data Availability

All code written to run the experiments and analyze the data presented here can be found at https://github.com/Neural-Control-Engineering/n-CORTex and https://github.com/Neural-Control-Engineering/AlphaModulation_SelectiveDetection. The data that support the findings of this study are available from the corresponding author upon reasonable request.

---

**Supplementary Figure 1:**
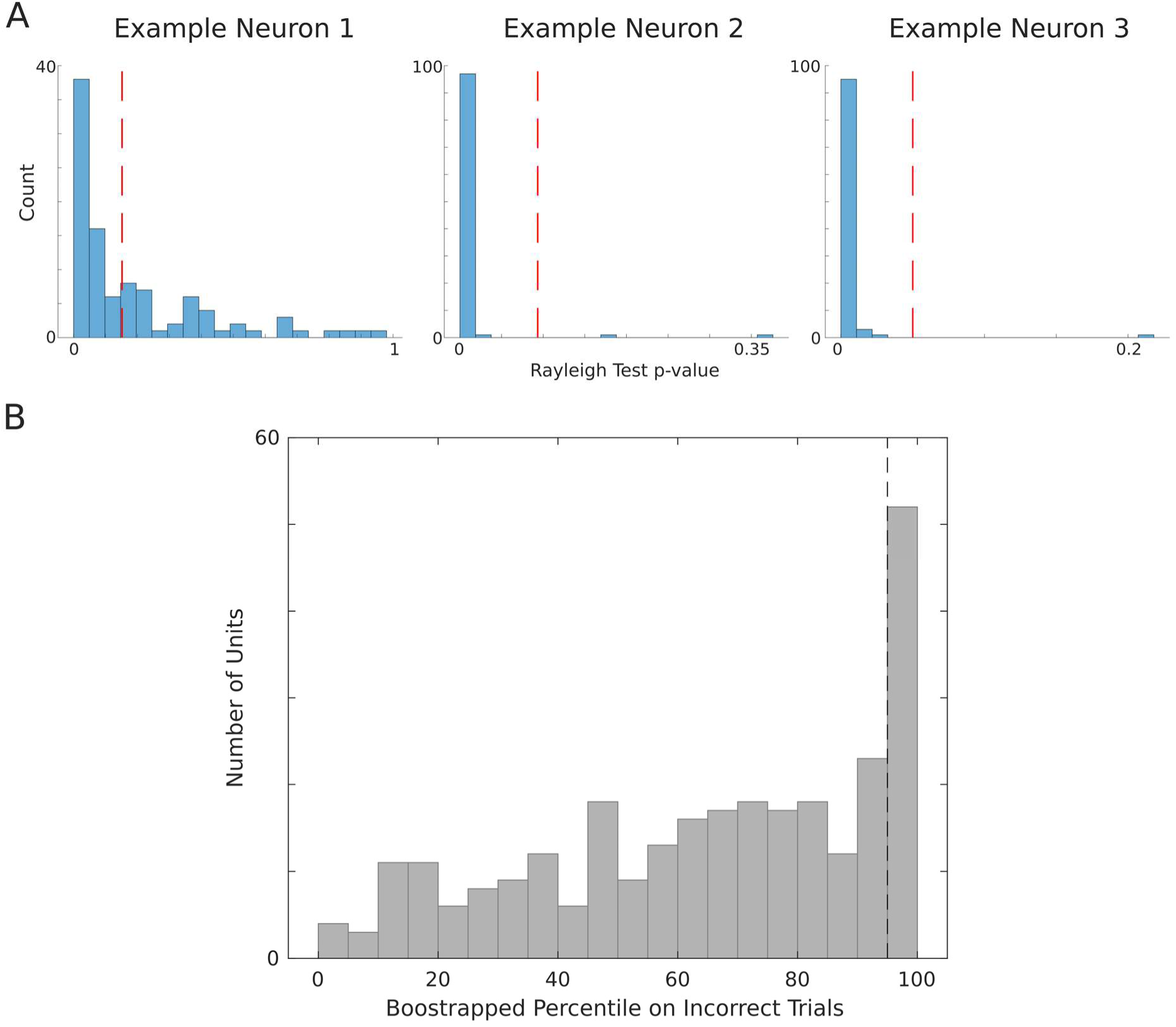
A bootstrapping method revealed that many phase modulated neurons exhibit significantly reduced phase modulation prior to incorrect trials. (A) Distributions of Rayleigh’s test p-values across 100 shuffles of correct trials matched to the number of incorrect trials (blue histograms) and the Rayleigh’s test p-value for across incorrect trials (red dashed lines) for three example neurons. (B) We computed the percentile of the Rayleigh’s test p-value across incorrect trials with respect to the distribution of p-values across correct trial-matched shuffles for all neurons that were selectively phase modulated prior to correct trials. Black dashed line indicates 95^th^ percentile.

**Supplementary Figure 2:**
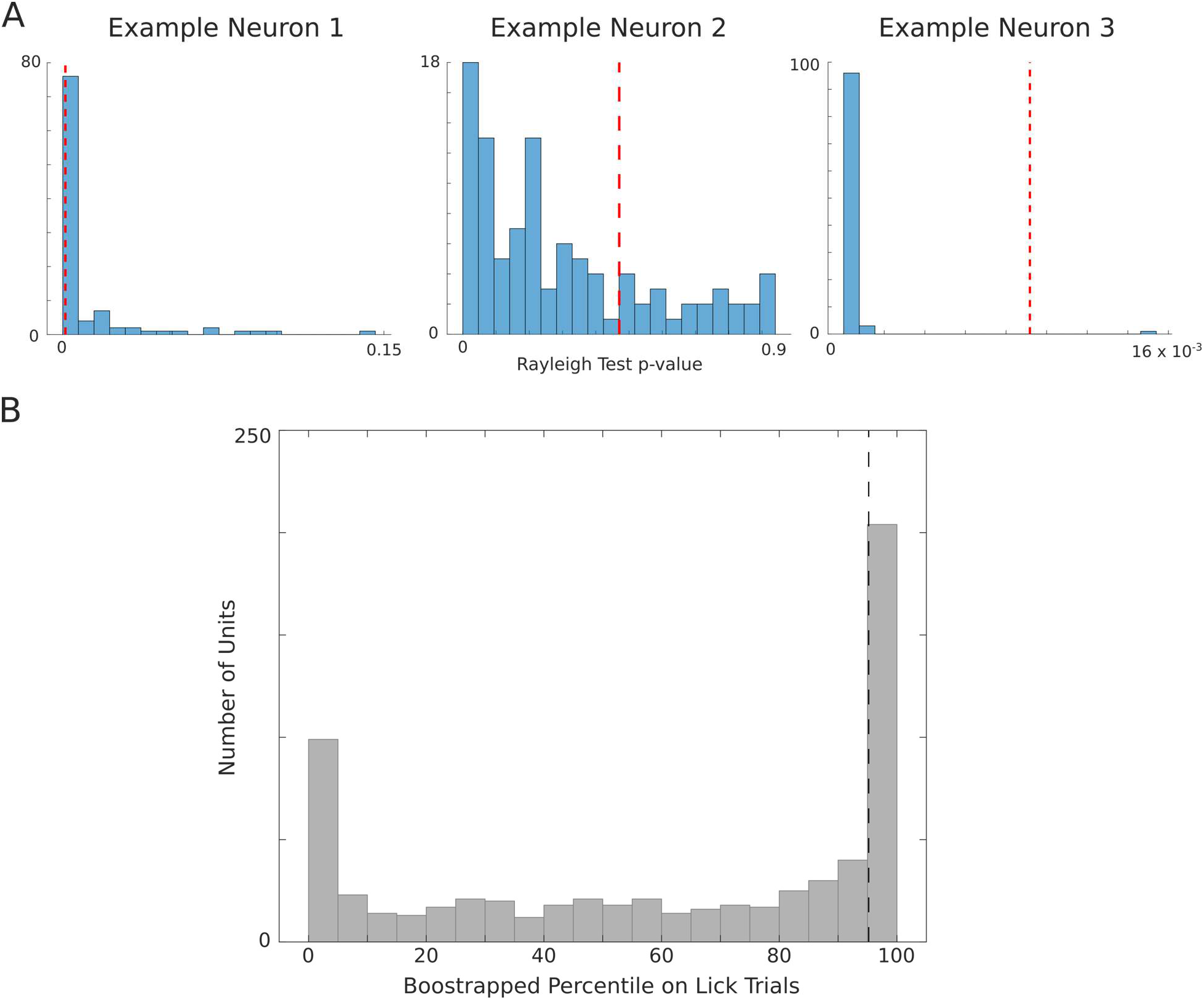
A bootstrapping method revealed that many phase modulated neurons exhibit significantly increased and decreased phase modulation during inter-trial intervals containing spontaneous licking. (A) Distributions of Rayleigh’s test p-values across 100 shuffles of trials without spontaneous licks matched to the number of trials with spontaneous licks (blue histograms) and the Rayleigh’s test p-value for across trials with spontaneous licks (red dashed lines) for three example neurons. (B) We computed the percentile of the Rayleigh’s test p-value across trials with spontaneous licks with respect to the distribution of p-values across no-lick trial-matched shuffles for all phase modulated neurons. Black dashed line indicates 95^th^ percentile.

## Notes

### Summary of Updates

This version of the manuscript has been revised to include many new analyses.

